# Diverse communities behave like typical random ecosystems

**DOI:** 10.1101/596551

**Authors:** Wenping Cui, Robert Marsland, Pankaj Mehta

## Abstract

With a brief letter to *Nature* in 1972, Robert May triggered a worldwide research program in theoretical ecology and complex systems that continues to this day[1]. Building on powerful mathematical results about large random matrices, he argued that systems with sufficiently large numbers of interacting components are generically unstable. In the ecological context, May’s thesis directly contradicted the longstanding ecological intuition that diversity promotes stability[2–4]. In economics and finance, May’s work helped to consolidate growing concerns about the fragility of an increasingly interconnected global marketplace[5–7]. In this Letter, we draw on recent theoretical progress in random matrix theory and statistical physics to fundamentally extend and reinterpret May’s theorem. We confirm that a wide range of ecological models become unstable at the point predicted by May, even when the models do not strictly follow his assumptions. Surprisingly, increasing the interaction strength or diversity beyond the May threshold results in a reorganization of the ecosystem – through extinction of a fixed fraction of species – into a new stable state whose properties are well described by purely random interactions. This self-organized state remains stable for arbitrarily large ecosystem and suggests a new interpretation of May’s original conclusions: when interacting complex systems with many components become sufficiently large, they will generically undergo a transition to a “typical” self-organized, stable state.

---

For an ecosystem of *S* species, May’s theorem concerns the *S* × *S* community matrix **J**, whose entries *J*_*ij*_ describe how much the growth rate of species *i* is affected by a small change in the population *N*_*j*_ of species *j* from its equilibrium value 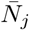 [1]. The stability of this equilibrium can be quantified in terms of the largest eigenvalue *λ*_max_ of **J**. If *λ*_max_ is positive, the equilibrium is unstable, and a small perturbation will cause the system to flow away from the equilibrium state. In the 1950’s, Eugene Wigner derived a mathematical formula for the distribution of eigenvalues in a special class of large random matrices[8]. May pointed out that the resulting estimate of the maximum eigenvalue *λ*_max_ is actually more general, and applies whenever the *J*_*ij*_ are sampled independently from probability distributions with finite means *µ* and variances *σ*^2^[1]. With this result in mind, May considered a simple ecosystem where each species inhibits itself, with *J*_*ii*_ = −1, but different species initially do not interact with each other. This ecosystem is guaranteed to be stable for any level of diversity. He then examined how the stability is affected by adding randomly sampled interactions, and found that *λ*_max_ typically becomes positive when the root-mean-squared total strength 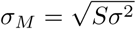 of inter-specific interactions reaches parity with the intra-specific interactions, that is, when *σ*_*M*_ = 1. For a given pairwise interaction strength *σ*, this relation gives the maximum diversity *S* compatible with ecosystem stability.

This result has proven to be robust against a wide array of changes in the assumptions, including adding biologically realistic correlation structures to the matrix, or incorporating the dependence of the community matrix on population sizes in the Lotka-Volterra model [9, 10]. In order to be stable at high diversities, a generic ecosystem must have a fine-tuned interaction structure, which can sometimes be justified in terms of biological constraints[11–14]. Very recently, it was noted that the required level of structure may emerge spontaneously in simple models of population dynamics, typically through extinction of some fraction of the initial set of species [15, 16]. Surprisingly, the resulting self-organized states were relatively insensitive to the network structure encoded in the interaction matrix. These works suggest that May’s result might admit of a more intuitive interpretation, and that the high-diversity states likely possess interesting generic properties.

To explore these ideas, we devised a more concrete version of May’s original thought experiment describing an ecosystem consisting of *S* non-interacting species where interactions are gradually turned on. May’s original argument only considered the local dynamics near a prespecified equilibrium point that eventually becomes unstable. Since we are interested in exploring what happens after the onset of this instability, we must make additional modeling assumptions to arrive at a complete set of nonlinear dynamics. We focus on MacArthur’s Consumer Resource Model (CRM) where interactions are mediated by competition for *M* substitutable resources[16]. For simplicity, we assume that *S* = *M* (though this does not affect our main results). MacArthur’s model allows for a wide range of exact mathematical results, but assumes that the resources are themselves self-replicating entities. To check the generality of our results, we also numerically analyzed generalizations of the CRM including the case of constant flux of externally supplied resources, and a model of microbial ecology with trophic feedbacks where organisms can feed each other via metabolic byproducts[17, 18]. These extended results can be found in the Supplemental Information.

The identity of each species in these models is determined by its consumption preferences. A set of noninteracting species can be constructed by engineering each species to consume a different resource type, with no overlap between consumption preferences. One can imagine designing strains of *E*. *coli* where one strain only expresses transporters for lactose, and another only expresses transporters for sucrose, etc., with all other transporters edited out of the genome, as illustrated in Figure 1(A). In such an experiment, horizontal gene transfer would eventually begin distributing transporter genes from one strain to another, so a realistic model would have to allow for some amount of unintended resource consumption. The resulting preference 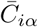 of species *i* for resource *α* is the sum of the identity matrix 1 and a random component *C*_*iα*_ with variance *σ*^2^.

**FIG. 1.**
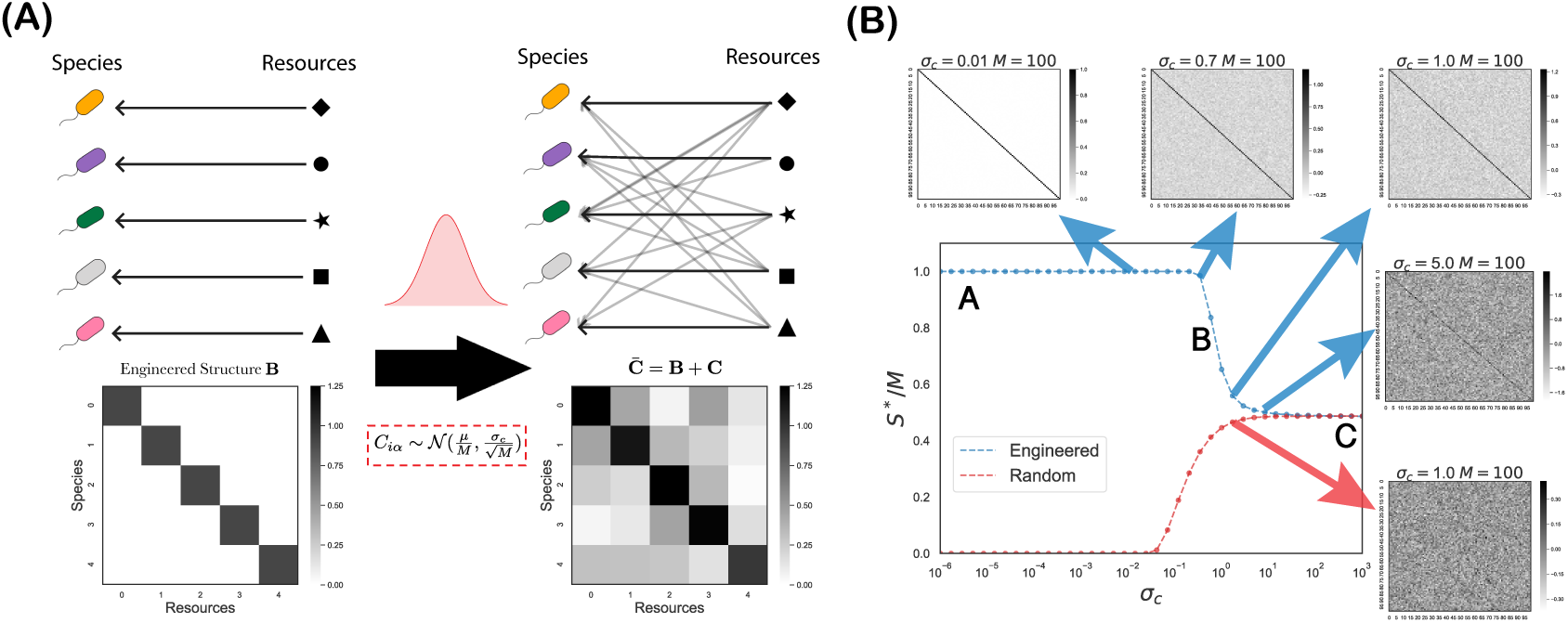
Random interactions destabilize an ecosystem of specialist consumers. **(A)** Left: an ecosystem with system size *M* = 5 starts with specialists consuming only one type of resource, resulting in a consumer preference matrix **B** = 1. Right: off-target consumption coefficients 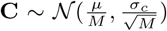 are sampled from a Gaussian distribution, resulting in an overall consumer preference matrix 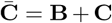. **(B)** Fraction of surviving species *S*\*/*M* vs. *σ*_*c*_, numerically computed using *M* = 100 as described in the Methods, along with the corresponding results for a completely random ecosystem with **B** = 0. Also shown are examples of the matrices 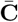 employed in the simulations.

Figure 1(B) shows the results of adding non-specific interactions to the CRM. Just as in May’s analysis, the appropriate measure of the importance of the random component is the root-mean-squared off-target consumption 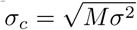 (recall *M* = *S*). Since we are expecting some species to go extinct in the self-organization process, we start by plotting the fraction of surviving species *S*\*/*M* versus *σ*_*c*_ in the steady state of numerical simulations with *S* = 100 species. At small values of *σ*_*c*_, all the species survive, as expected for a weakly interacting ecosystem. As high as *σ*_*c*_ = 0.7, almost all of the original species are still present in the community. But between *σ*_*c*_ = 0.7 and *σ*_*c*_ = 1, there is a sharp transition in community structure, which results in about half of the original species becoming extinct. Remarkably, the survival fraction converges to the same value as for a completely random consumer preference matrix, and remains finite as *σ*_*c*_ → ∞. This means that arbitrarily high amounts of diversity can be maintained at a given amount of uncertainty *σ* in the model parameters by considering a sufficiently large starting ecosystem. These numerical predictions are in excellent agreement with analytic predictions for the limit *S* → ∞ derived in the Methods.

We proceeded to investigate the properties of these self-organized high-diversity states more closely. In addition to the number of surviving species, we considered two other community-level properties: the mean population size ⟨*N*⟩ over all species, and the second moment of the population size ⟨*N*^2^⟩, which includes information about the range of population sizes. Figure 2 shows that both of these quantities are also well-approximated by the random consumer preference matrix for *σ*_*c*_ > 1. This convergence to random ecosystem behavior is quite robust, and remains present when a set of designed interactions is added to the original noninteracting community before adding the random component. Figure 2 shows the results for two basic interaction structures: a block structure with pre-defined groups of species exhibiting strong intra-group competition, and a unimodal structure where each species is more likely to consume resources similar to its preferred resource. The only effect of the choice of structure is to adjust the threshold value of *σ*_*c*_ where the transition takes place. The character of the self-organized state is also robust to changes in the sampling scheme for the random component. Gaussian sampling allows the clearest comparison to May’s result, but it produces some negative values, while consumer preferences should always be positive. We therefore tested two sampling schemes that always produce positive values: uniform sampling in the interval from 0 to *b*, and binary sampling with probability *p* of choosing 1. Changing *b* or *p* affects both the mean and the variance of the random component, and so the transition point cannot be directly compared to the Gaussian case. But we still find that the large-scale properties become similar to those of the random ecosystem when the average total off-target consumption capacity over all *M* resource types becomes greater than the original designed capacity.

**FIG. 2.**
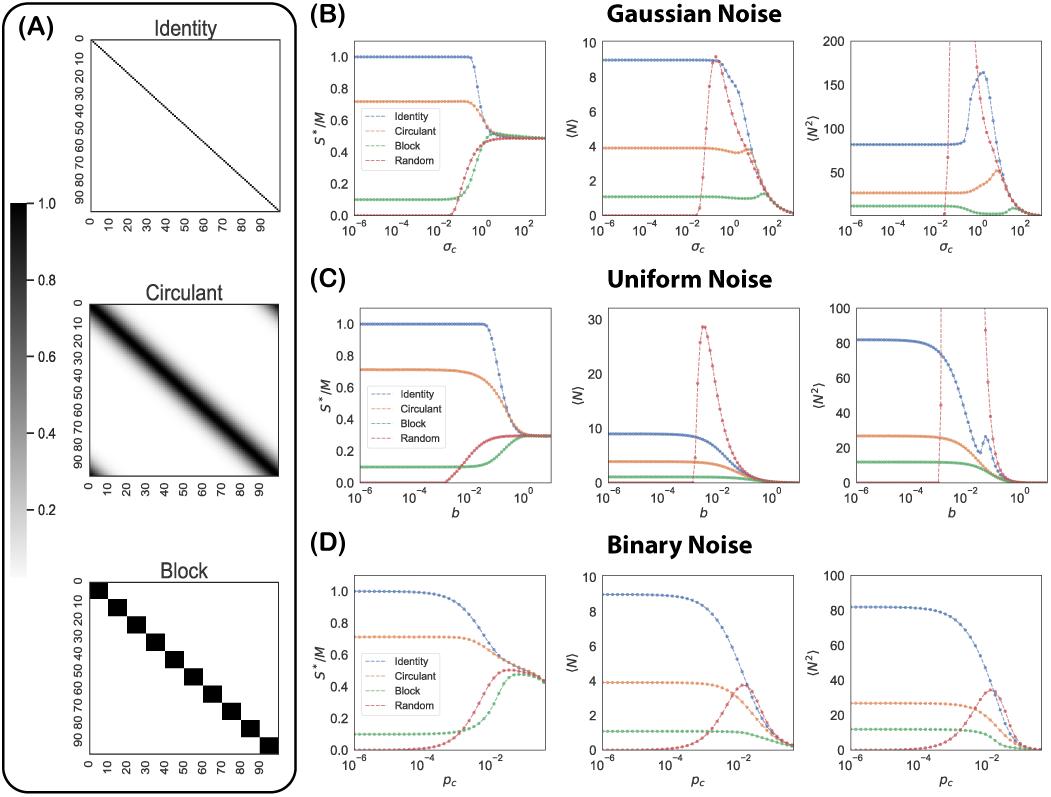
Community properties for structured and random ecosystems. **(A)**: Examples of designed interactions **B**. Top: an identity matrix; Middle: a Gaussian-type circulant matrix; Bottom: a block matrix. **(B)** Gaussian Noise 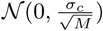, **(C)** Uniform Noise: 𝒰 (0, *b*) and **(D)**: Binary Noise: *Bernoulli*(*p*_*c*_) are added to the engineered interactions. Community properties including survival fraction *S*\*/*M*, mean abundance ⟨*N*⟩ and mean square abundance ⟨*N*^2^⟩ are shown for the different **B** listed above.

Finally, we examined the stability of the ecosystems. For comparison with May’s results, we obtained effective competition coefficients *A*_*ij*_ between species in a generalized Lotka-Volterra model, by assuming that the resource abundances always remain close to their steady state values, as illustrated in Figure 3(A). This matrix is related to May’s community matrix by 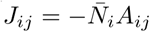. For the symmetric interaction matrices arising from the consumer resource model, one can prove that the largest eigenvalue *λ*_max_ of **J** reaches zero from below only when the smallest eigenvalue *λ*_min_ of **A** reaches zero from above (see Methods). Figure 3(B) shows how the eigenvalues of **A** change as *σ*_*c*_ increases. Initially, all the eigenvalues of the noninteracting community are identical, but with increasing *σ*_*c*_ the distribution spreads out, and reaches the threshold of stability *λ*_min_ = 0 at *σ*_*c*_≈ 1. In the Methods we show analytically using the Cavity method [20–21] that the transition point approaches 1 as *S* → ∞, in agreement with expectations based on May’s analysis.

**FIG. 3.**
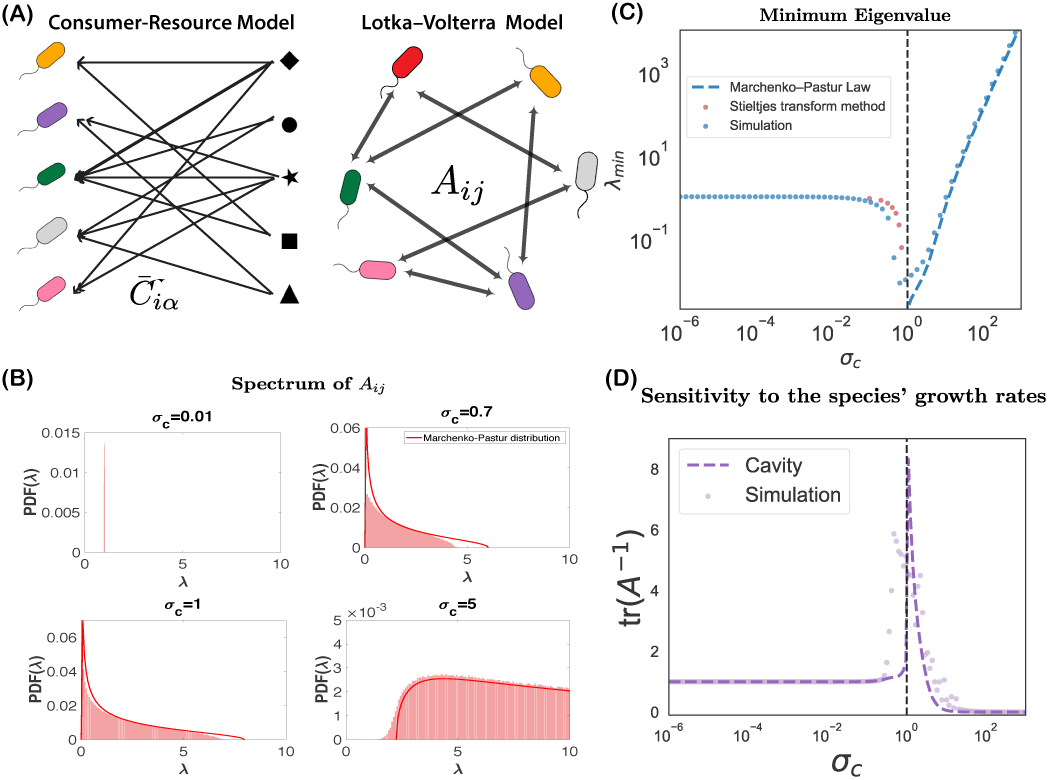
Effect of random interactions on ecosystem stability. **(A)**: The bipartite interactions 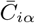 in MacArthur’s consumer-resource model can be mapped to pairwise competition coefficients *A*_*ij*_ in generalized LotkaVolterra equations through 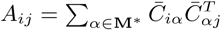. **(B)** Spectra of *A*_*ij*_ at different *σ*_*c*_ for **B** = 𝟙. The red solid line is the Marchenko-Pastur distribution. **(C)**: Comparison between numerical simulations and analytic results for the minimum eigenvalue of **A** at different *σ*_*c*_. **(D)**: Comparison between numerical simulations and analytic solutions for the mean sensitivity *v* of steady-state population sizes to changes in species’ growth rates. See Methods for detailed calculations.

But for *σ*_*c*_ > 1, we observe two new phenomena that were not accessible in May’s original framework. First, the spectrum is well-described by the eigenvalue distribution for interactions resulting from completely random consumer preference matrices, known as the Marchenko-Pastur Distribution[22]. This is consistent with our earlier observations on community-level observables that all coincided with the random case for sufficiently large *σ*_*c*_. Secondly, the minimum eigenvalue in the Marchenko-Pastur Distribution is located at 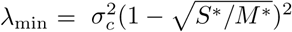, where *M\** is the number of resource types that persist at nonzero abundance in the steady state. As we saw earlier, about half the species go extinct when *σ*_*c*_ > 1, leading to *S*\*/*M*\* < 1, so that *λ*_min_ is always positive. This means that the random ecosystem is stable, for arbitrarily large values of *σ*_*c*_.

The spectrum of **A** also contains quantitative information about the degree of ecosystem stability. Specifically, as shown in the Methods, the sum of the inverse eigenvalues Σ _*i*_ (1*/λ*_*i*_) = tr(**A**^*-*1^) measures the average response of the steady-state population size 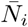 to a given perturbation of the species’ growth rate. Figure 3(D) shows that this quantity is initially constant as *σ*_*c*_ is increased from 0, then diverges at *σ*_*c*_ = 1, and finally rapidly decreases to near zero. In the Methods we provide analytical calculations based on the cavity method confirming that this divergence is a signature of a continuous phase transition, analogous to the increased sensitivity near ecological tipping points known to result in large stochastic fluctuations [23, 24].

The foregoing analysis leads to a reinterpretation of May’s theorem as a bound on the feasibility of bottom-up engineering in complex systems. As the number of components increases, small uncertainties in each of the interaction parameters eventually overwhelm the designed interactions, and destabilize the intended system state. But the system generically finds a new typical stable state which may be even more stable than the originally engineered one. Importantly, our work suggests that crossing the May transition generically gives rise to typical random ecosystems rather than a specialized phase as was found in a recent analysis of the Generalized Lotka-Volterra model [16]. For this reason even when the cumulative parameter uncertainties preclude a priori prediction of the detailed structure of the new state, methods from statistical physics and Random Matrix Theory can be employed to predict system-level properties [15, 25]. Further development of these methods and their applications will play an important role in enabling top-down control of systems beyond the May bound and helping to identify assembly rules for microbial communities [26]

## ACKNOWLEDGMENTS

We thank Zhenyu Liao, Guangwei Si, Jean Vila and Yu Hu for helpful discussions. The work was supported by NIH NIGMS grant 1R35GM119461, Simons Investigator in the Mathematical Modeling of Living Systems (MMLS). The authors are pleased to acknowledge that the computational work reported on in this paper was performed on the Shared Computing Cluster which is administered by Boston University Research Computing Services.

## METHODS

### A. Model

In this work, we will focus primarily on MacArthur’s consumer-resource model(MCRM)[17]. This model consists of *S* species or consumers with abundances *N*_*i*_ (*i* = 1…*S*) that can consume one of *M* substitutable resources with abundances *R*_*α*_ (*α* = 1…*M*)

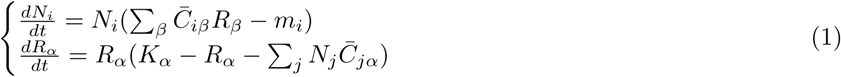

The consumption rate of species *i* for resource *α* is encoded by the element 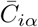 in the *S* × *M* matrix 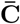. *K*_*α*_ is the carrying capacity of each resource *α*. *m*_*i*_ is some minimum maintenance cost that species *i* must harvest from resources in order to grow. Note, both the species and resource abundances *N*_*i*_ and *R*_*α*_ must be strictly non-negative. When the system is in the steady state, some species and resources can vanish. We denote the numbers of surviving species and resources by *S*\* and *M* *, respectively.

To construct our mechanistic version of May’s analysis, we decompose the consumer matrix 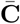 into two parts: 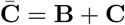 with **B** encoding a pre-designed set of resource-mediated interactions, and **C** a random matrix encoding “off-target” consumption. We consider three types of **B** (see Figure 2): the identity matrix, a square Gaussian-type circulant matrix 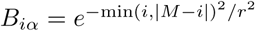 with *r* = 7[27] and a block matrix with identical 10 × 10 blocks(all elements are 1 inside the 10 × 10 block). We also consider three types of random matrices **C**. In all cases, each element in the matrix is sampled independently from an underlying probability distribution. The three distributions we consider are a normal distribution with mean zero and standard deviation 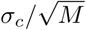, a uniform distribution where each element is sampled uniformly from [0, *b*], and a Bernoulli distribution where each element can be +1 with probability *p*_*c*_ and 0 with probability 1 *- p*_*c*_ (i.e Binary Noise).

For all simulations, unless otherwise specified the default choices for parameters are: *M* = 100, *µ* = 0, *K* = 10, *σ*_*K*_ = 1, *m* = 1 and *σ*_*m*_ = 0.1 and each data point is averaged from 4000 independent realizations. For Figure 3 (C, D), each data point is averaged from 8000 independent realizations. All simulations are available on GitHub at https://github.com/Emergent-Behaviors-in-Biology/typical-random-ecosystems.

#### 1. Alternative Models used in SI

To test the generality of our results, we also simulated more complicated variants of the consumer resource model (see Figure S1). First, we simulated a consumer resource model with linear resource dynamics. In this case, the second equation in (1) is replaced by

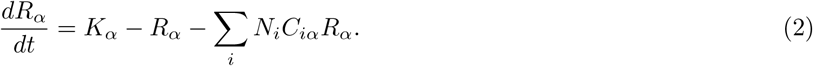

This small change can significantly change the ecosystem properties, because it prevents resources from going extinct in the steady state. In the simulations, we set *M* = 100, *µ* = 1, *K* = 10, *σ*_*K*_ = 1, *m* = 1 and *σ*_*m*_ = 0.1 and each data point is averaged from 4000 independent realizations.

Second, we simulated a generalization of the MacArthur’s Consumer Resource model we call the Microbial Consumer Resource Model (MicroCRM). The MicroCRM was introduced in [18] and refined in [19] to simulate microbial communities. In this model, in addition to consuming resources species can produce new resources through crossfeeding. This dramatically changes the resource dynamics through the introduction of trophic feedbacks. Unlike the original CRM and the extensio to linear resource dynamics, the MicroCRM possesses no Lyapunov function. Full details of the model are available in the appendix of [19]. In particular, the dynamics we use are described in equation (17)[19] with the leakage rate *l* = 0.4. The fraction of secretion flux secreted to the same resource type is *f*_*s*_ = 0.45, the fraction of secretion flux to ‘waste’ resource is *f*_*w*_ = 0.45 and variability in secretion fluxes among resources is *d*_0_ = 0.2. We set *M* = 100, *µ* = 1, *K* = 10, *σ*_*K*_ = 1, *m* = 1 and *σ*_*m*_ = 0.1 and each data point is averaged from 4000 independent realizations.

### B. Sensitivity to Parameter Perturbations

We begin by defining four susceptibility matrices that measure how the steady-state resource and species abundances respond to changes in the resource supply and species death(growth) rates:

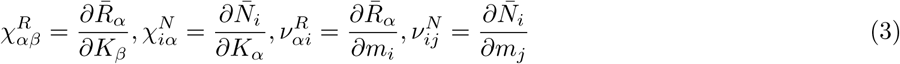

where the bar 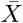 over the variable *X* denotes the steady-state (equilibrium) solution.

For the extinct species and resources, by definition the susceptibilities are zero. For this reason, we focus only on the surviving resources and species. At steady-state, equation (1) gives:

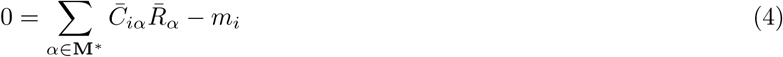

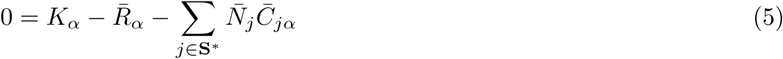

where **M***\** and **S***\** denote the sets of resources and species, respectively, that survive in the ecosystem at steady-state. Differentiating these equations yields the relations

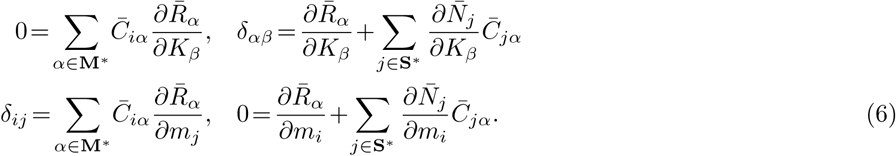

Substituting in for the partial derivatives using the susceptibility matrices defined above, we have:

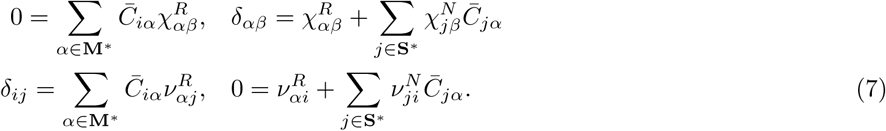

These two equations can be written as single matrix equation for block matrices:

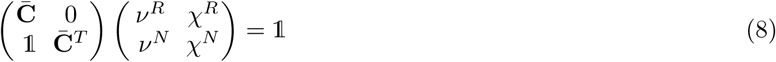

To solve this equation, we define a *S*\* × *S*\* matrix: 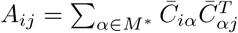. A straightforward calculation yields

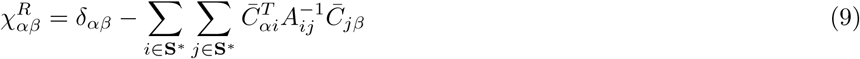

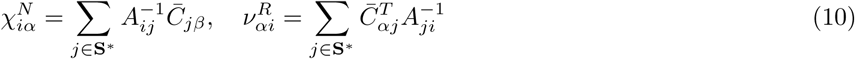

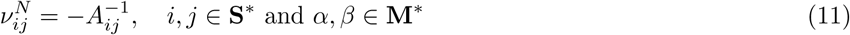

### C. Cavity Solution

For an initially non-interacting ecosystem **B** = 𝟙, the effect of random off-target consumption on system-scale properties can be computed analytically in the *M, S* → ∞ limit using the cavity method [20, 21]. The cavity calculation is straightforward but tedious. For this reason, it is helpful to introduce the notation:

- 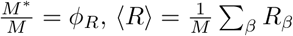 and 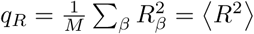, where *M* * is the number of surviving resources.
- 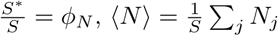 and 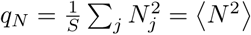, where *S*\* is the number of surviving species.
- 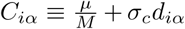 assuming 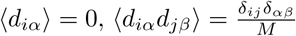. with 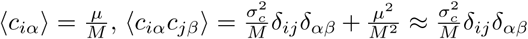.
- *K*_*α*_ = *K* + *dK*_*α*_ with 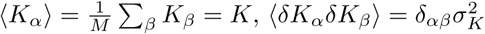.
- *m*_*i*_ = *m* + *dm*_*i*_ with 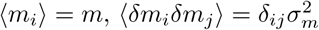.
- 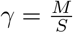 and for the identity matrix *γ* = 1.

Following similar steps as in [21], we perturb the ecosystem with a new species and resource *N*_0_ and *R*_0_. Ignoring 𝒪(1*/M*) terms yields the following equations:

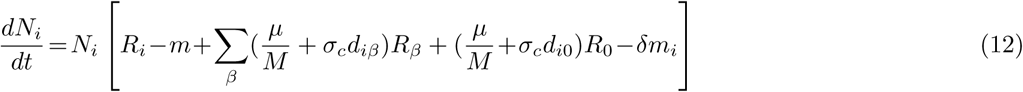

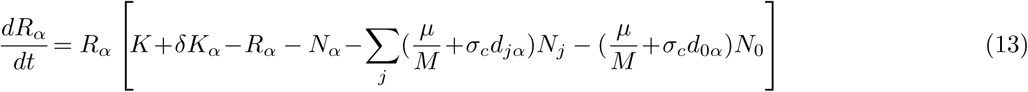

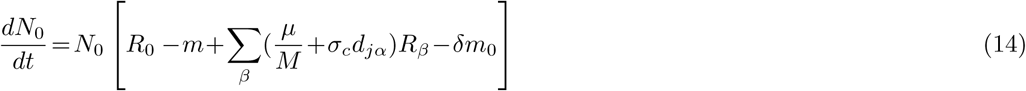

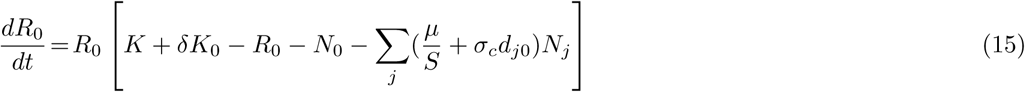

Denote by 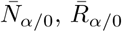 and 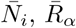 the equilibrium values of the species and resources before and after adding the newcomers, respectively. These can be related to each other using the susceptibilities defined above:

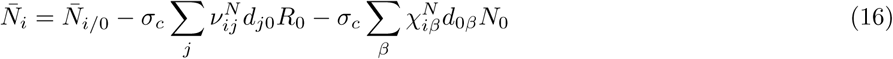

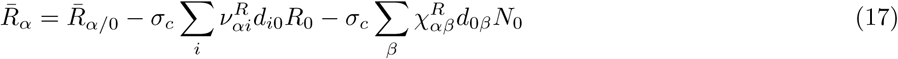

It is is helpful to introduce new auxiliary random variables:

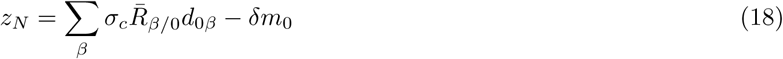

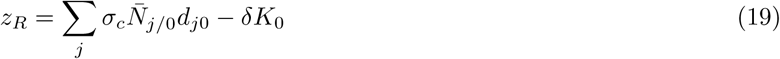

where 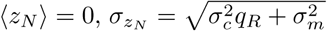 and 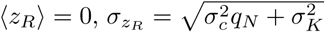 Following calculations analogous to [21] and noting that 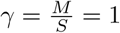 yields:

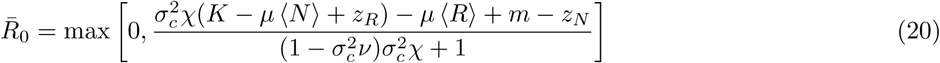

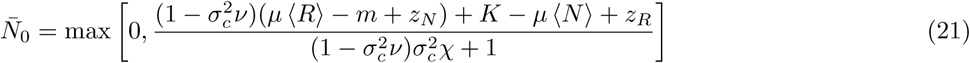

Cavity equations for the susceptibilities can be obtained directly by differentiating these equations:

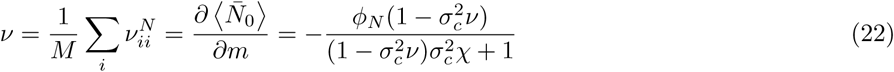

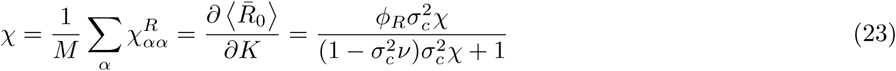

Two solutions are found:

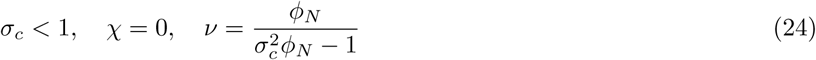

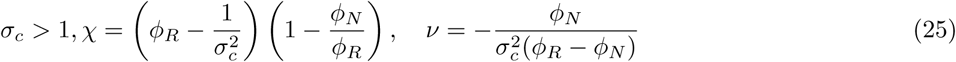

The comparison between cavity solutions and numerical simulations are given in Figure S3 and Figure 3(C).

#### 1. Three Regimes of Behavior

To understand these solutions and behaviors better, it is helpful to consider three regimes: Regime A where *σ*_*c*_ ≪ 1, Regime B where *σ*_*c*_ ≈ 1, and Regime C where *σ*_*c*_ ≫ 1. Equation (25) shows the linear response function *χ* in Regime B: *σ*_*c*_ ∼ 1 and Regime C: *σ*_*c*_ ≫ 1 are only different at order 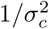. After the phase transition 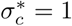, a slight increase of *σ*_*c*_ will induce a transition from Regime B into Regime C. This explains the dramatic drop of the species packing shown in Figure 1(B). In Regime A (*σ*_*c*_ ≪ 1), the equations for the steady-states become

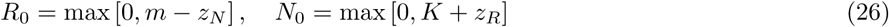

For Regime C (*σ*_*c*_ ≫ 1), the solution is well approximated by

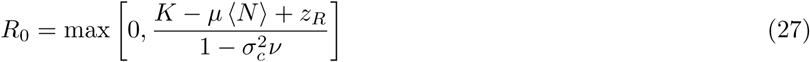

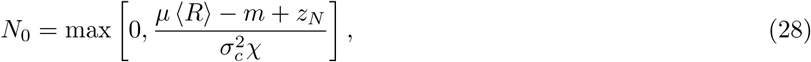

in agreement with the equations obtained in [21] for purely random interactions.

### D. Lotka-Volterra Model, Wishart Matrix and Marchenko-Pastur Law

In this section, we show how the generalized Lotka-Volterra model can be related to the CRM, and in particular, the how the steady states of the two models can be made to coincide. Solving for the steady-state values of the non-extinct resources by setting the bottom equation in (1) equal to zero gives:

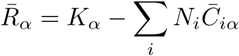

Substituting this into the top equation in (1) gives:

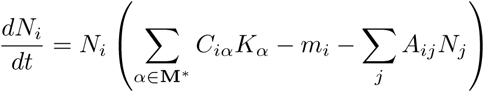

where we have defined an interaction matrix 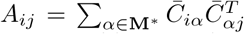 and **M**\* is the set of surviving resources. We can use this equation to solve for the steady-state (equilibrium) abundances of non-extinct species, and arrive at the expression:

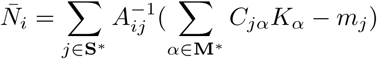

where **S**\* is the set of surviving species. In terms of 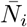, the Lotka-Volterra equations become:

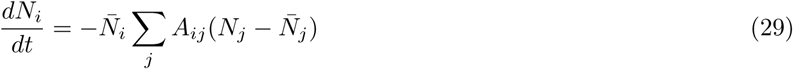

with community matrix

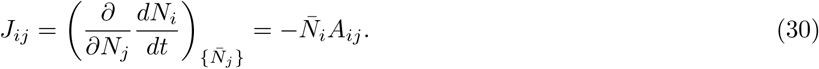

In May’s work, *J*_*i j*_ is assumed to be an i.i.d. random matrix and an extension of Wigner’s arguments about Gaussian random matrices is used to compute the leading eigenvalue [1]. Since the 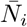 are not known *a priori*, the stability of Lotka-Volterra type dynamics are more easily studied in terms of the eigenvalues of *A*_*ij*_, using the connection between the leading eigenvalues of **J** and **A** derived below. Furthermore, for the Lotka-Volterra model obtained from the MCRM, **A** is the outer product of a random matrix 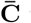 with itself, i.e., a Wishart matrix. The underlying reason for this is the bipartite nature of the MCRM resulting from the presence of two types of degrees of freedom: resources and species [28, 29]. Wishart matrices are well-known to follow a different eigenvalue distribution, the Marchenko-Pastur law[22] given by

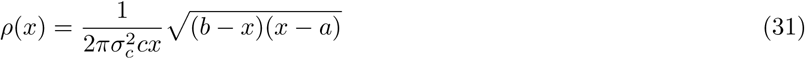

where 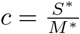.

For the problem we are studying, the bounds of the Marchenko-Pastur distribution are given by: 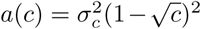 and 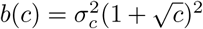. In Regime C: the random phase, the spectrum of **A** is extremely well described by *ρ*(*x*) (see Figure 3(B) and Figure S2). The Marchenko-Pastur law also allows us to find a analytic expression for the minimum eigenvalue of **A**:

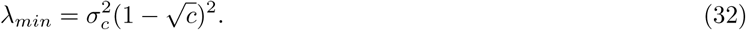

### E. Relating the eigenvalues of A and J

In this section, we prove that the largest eigenvalue *λ*_max_ of the community matrix **J** (which controls the Lyapunov stability of the fixed point) is negative if and only if the smallest eigenvalue *λ*_min_ of the Lotka-Volterra competition matrix **A** is positive. For this stability analysis, we remove the rows and columns corresponding to species that go extinct in the steady state, since allowing *N*_*i*_ = 0 trivially generates zero eigenvalues. **J** and **A** will always refer to the resulting matrices of dimension *S*\* × *S*\*.

We start by defining the diagonal matrix 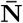, whose nonzero elements are the equilibrium population sizes 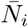. This lets us write

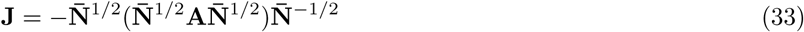

where 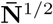 is the diagonal matrix whose entries are the square roots of the population sizes. This equation says that **J** is similar to 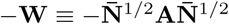, which implies that they share the same eigenvalues.

Since **W** and **A** are both symmetric matrices, their eigenvalues are all real, and the positivity of all the eigenvalues is equivalent to the positive-definiteness of the matrix.

Now we note that **W** is positive definite if and only if **A** is positive definite. For if **A** is positive definite, then **x**^*T*^ **Ax** > 0 for all column vectors **x** 0, including the column vector 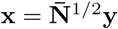 for any column vector **y** ≠ 0. But this implies that 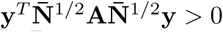 for all **y** ≠ 0, i.e., that **W** is positive definite. Conversely, if **W** is positive definite, then 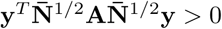 for all **y** ≠ 0, including 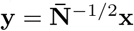 for any **x** ≠ 0. But this implies that **x**^*T*^ **Ax** > 0 for all **x** ≠ 0, i.e., that **A** is positive definite.

We conclude that the eigenvalues of **W** are all positive if and only if the eigenvalues of **A** are all positive. Therefore the largest eigenvalue of 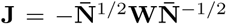 is negative if and only if the smallest eigenvalue of **A** is positive, as claimed in the main text.

### F. Correspondence between RMT and cavity solution

Our numerical simulations show that after the transition, our ecosystems are well described by purely random interactions. This suggests that we should be able to derive our cavity results using Random Matrix Theory (RMT). We now show that this is indeed the case. Our starting point are the average susceptibilities which are defined as:

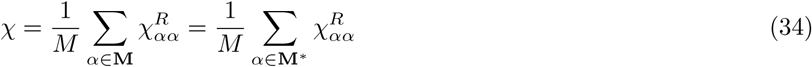

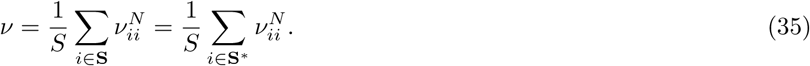

From the cavity calculations, we only care about 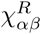 and 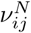, because the other susceptibilities are lower order in 1/*M*.

We can combine these equations with (10) and (11) to obtain

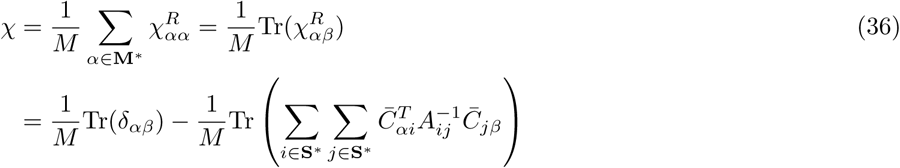

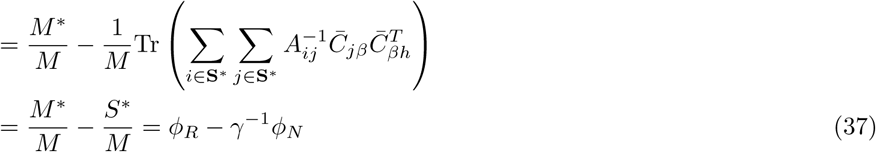

We now show that the cavity solutions are consistent with results from RMT using equations (10) and (11) in Regime A and Regime C described in the main text.

#### 1. Regime A: 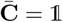

This regime happens when *σ*_*c*_ ≪ 1. Substituting, 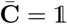 into equations (10) and (11) yields

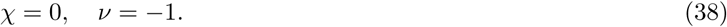

This is consistent with the cavity solution equation (24) with *σ*_*c*_ = 0 since in this case *S*\* = *S* = *M*.

#### 2. Regime C: 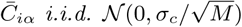

In this regime, *σ*_*c*_ ≫ 1. In this case, 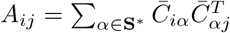 takes the form of a Wishart Matrix. We will exploit this to calculate 𝒳 and *v*. Notice,

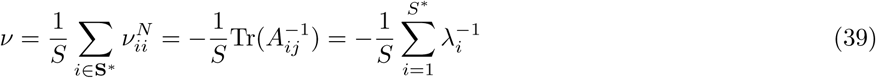

where *λ*_*i*_ is the eigenvalue of *A*_*ij*_. From the Marchenko-Pastur law [22], we know that the eigenvalues of a random Wishart matrix obey the Marchenko-Pastur distribution. Substituting equation (32) into the expression for *v* and replacing the sum with an integral yields:

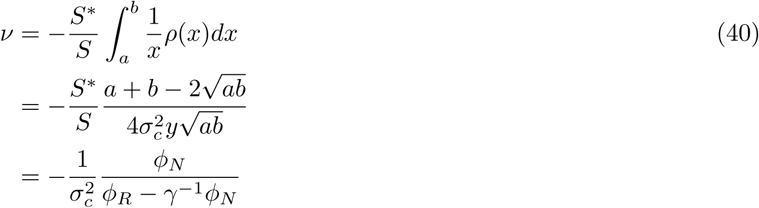

The second line of equation (40) is obtained by transferring the integral function to a complex analytic function and applying the residue theorem. This result is the same as the cavity solution equation (25) when *σ*_*c*_ ≫ 1.

#### 3. Regime B using the Stieltjes transformation

In Regime B, it hard to estimate the minimum eigenvalue. We can use Stieltjes transformation of information-plusnoise-type matrices which are well studied in wireless communications[30, 31], where **B** represents the information encoded in the signal and **C** is the noise in wireless communications. In this case, we have

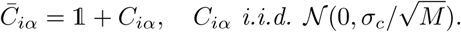

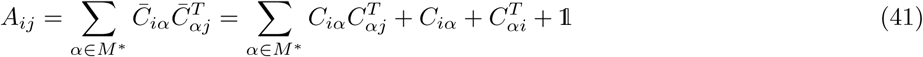

Using **Theorem 1.1** in Dozier and Silverstein[30], the Stieltjes transform *m*(*z*) of *A*_*ij*_ satisfies

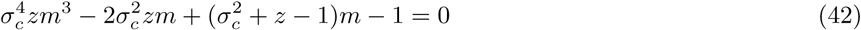

The asymptotic spectrum of *A*_*ij*_ can be obtained by *m*(*z*), the solution of equation (42) with

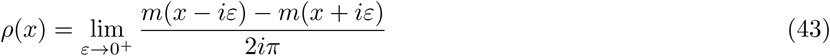

The result is shown in Figure S4. The minimum eigenvalue reaches 0 nearly at 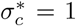, as predicted by the cavity solution.

## Supplementary Information

**FIG. S1.**
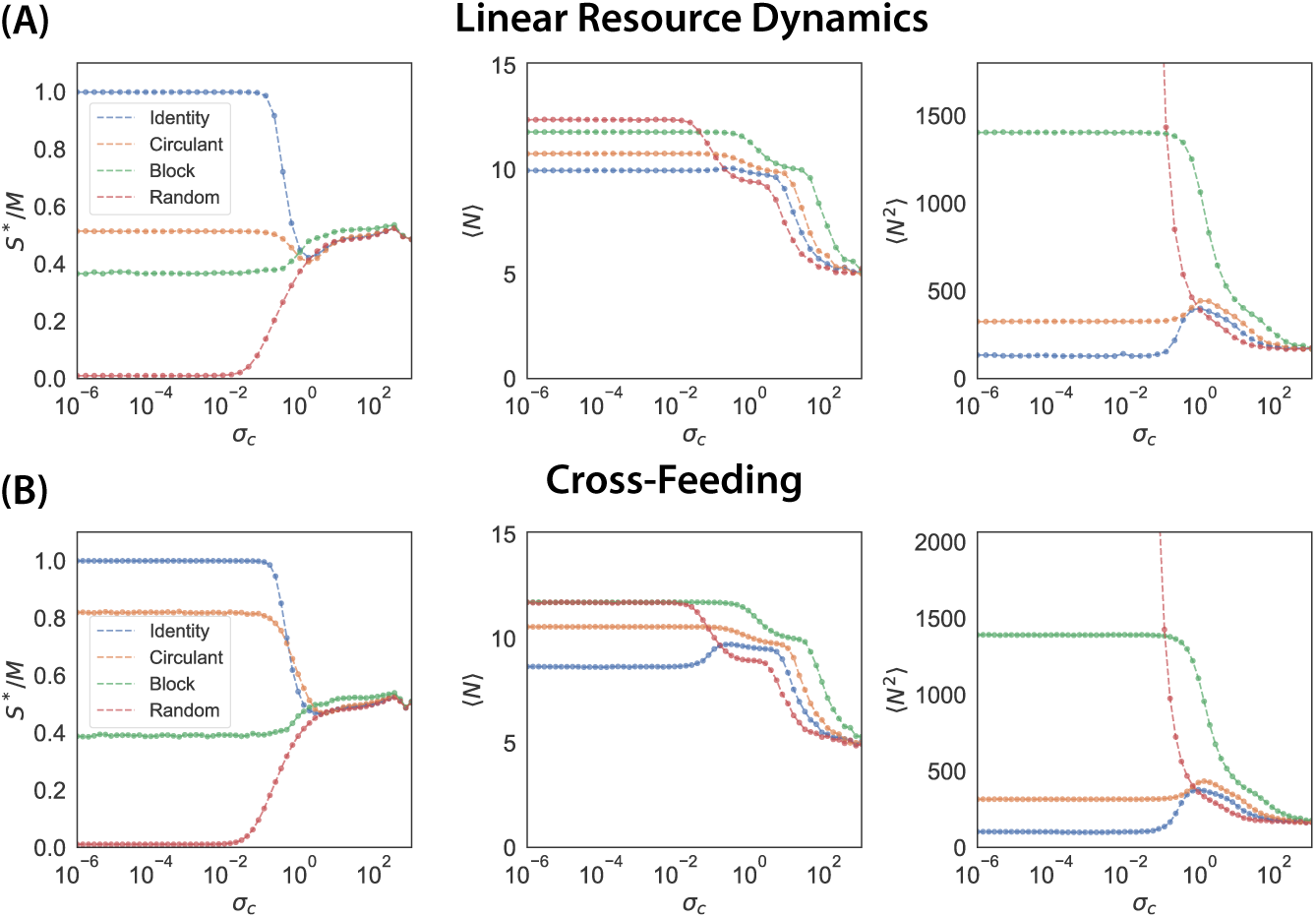
Community properties for generalized consumer-resource models under Gaussian noise. **(A)** Linear resource dynamics: the resource dynamics is changed to 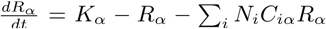. **(B)** With cross-feeding: the dynamics is described in equation (17) in Supplementary Information of [19]. The noise is only applied on the consumption matrix and *D* is kept the same at different *σ* _*c*_. In both models, **B** = 𝟙. See Methods for parameter values. Each data point is averaged over 4000 independent realizations.

**FIG. S2.**
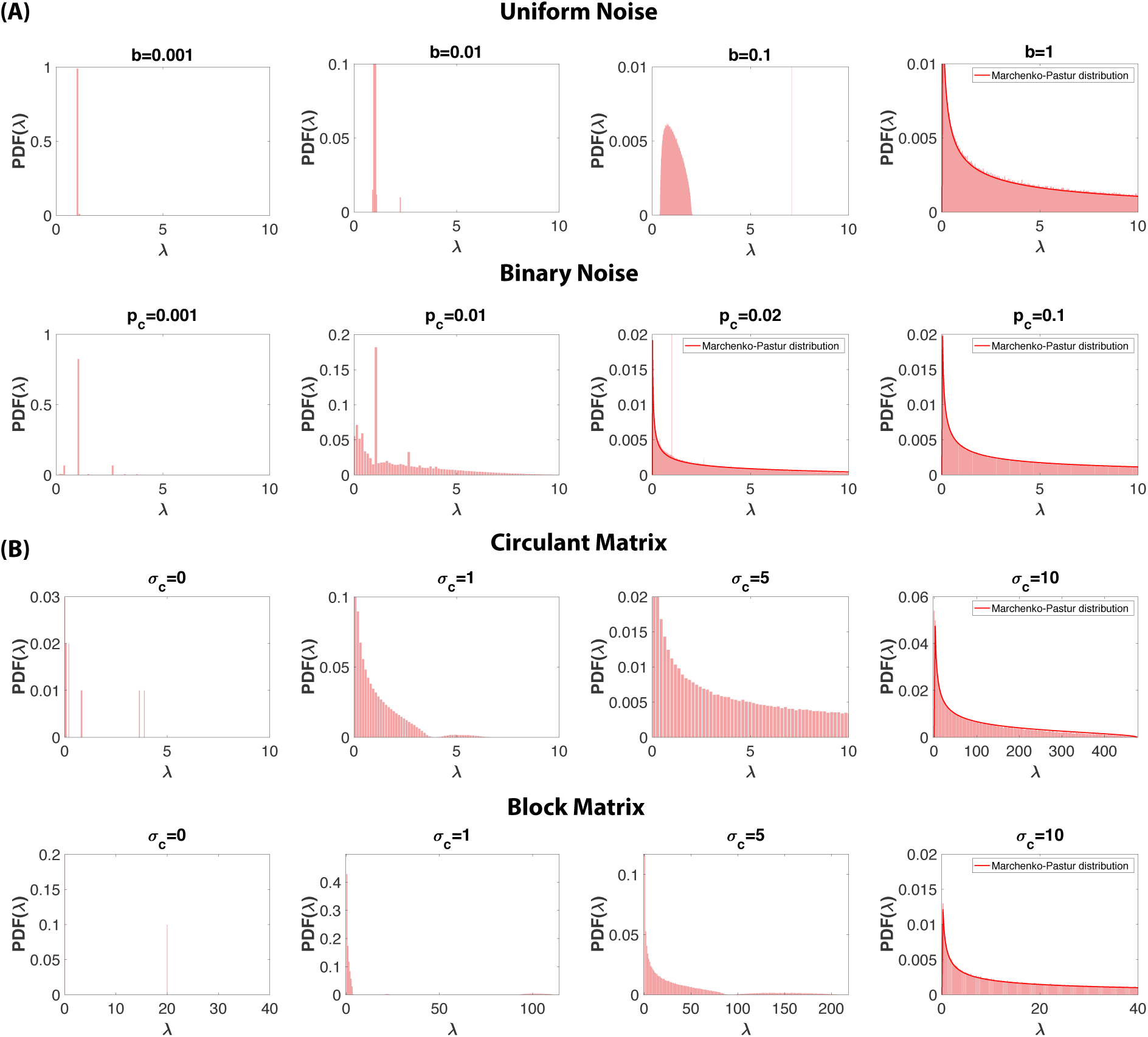
Spectra of *A*_*ij*_ in different cases. **(A)** Uniform Noise: 𝒰 (0, *b*) and Binary Noise: *Bernoulli*(*p*_*c*_); **B** is an identity matrix. **(B)** Gaussian noise and **B** is a circulant matrix. **(C)** Gaussian noise and **B** is a block matrix. Note that *S*,M \** in *A*_*ij*_ are obtained from numerical simulations.

**FIG. S3.**
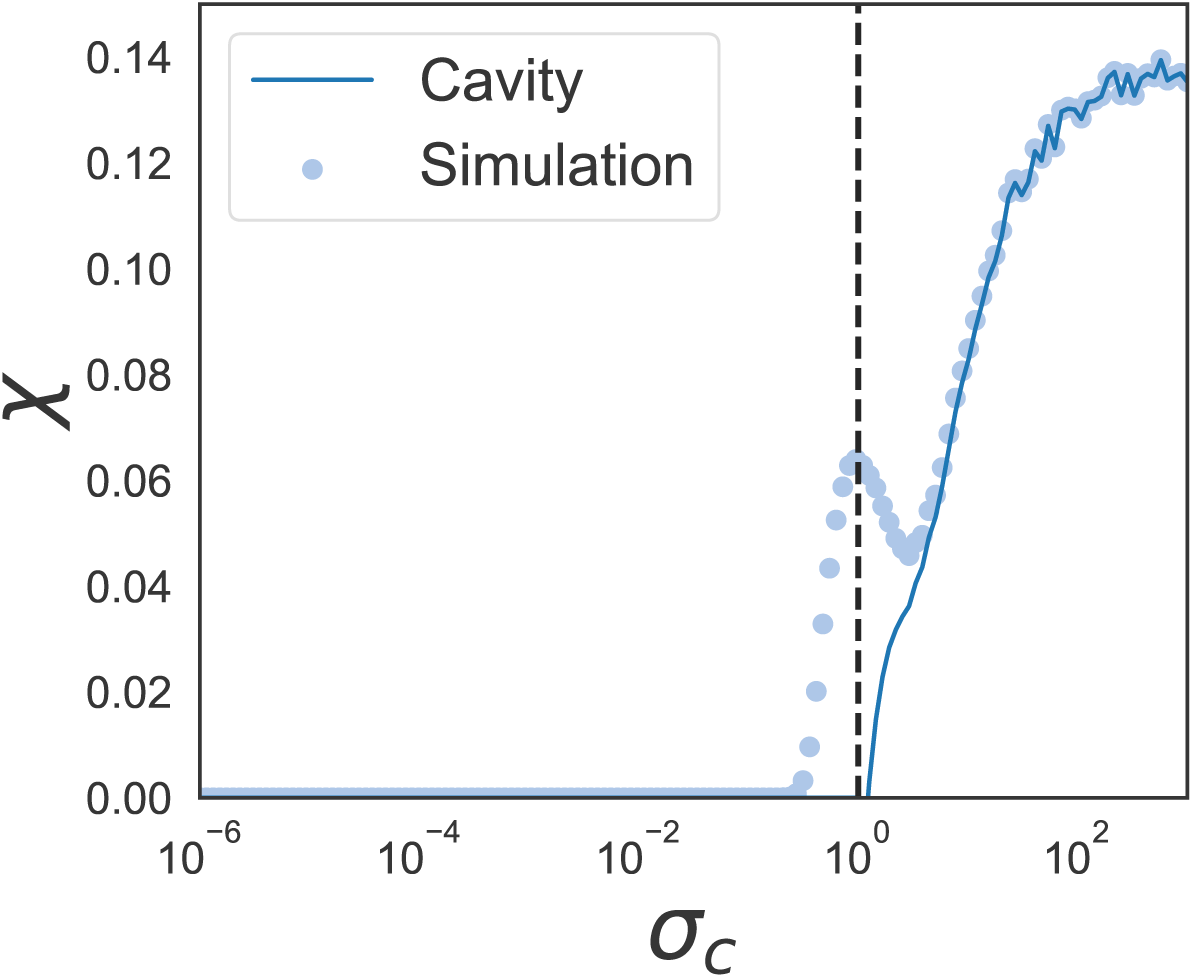
Comparison between numerical simulations and cavity solutions for *χ* at different *σ*_*c*_. Note *S\** and *M\** are obtained from the numerical simulations, although in principle they could be obtained by solving the cavity equations directly.

**FIG. S4.**
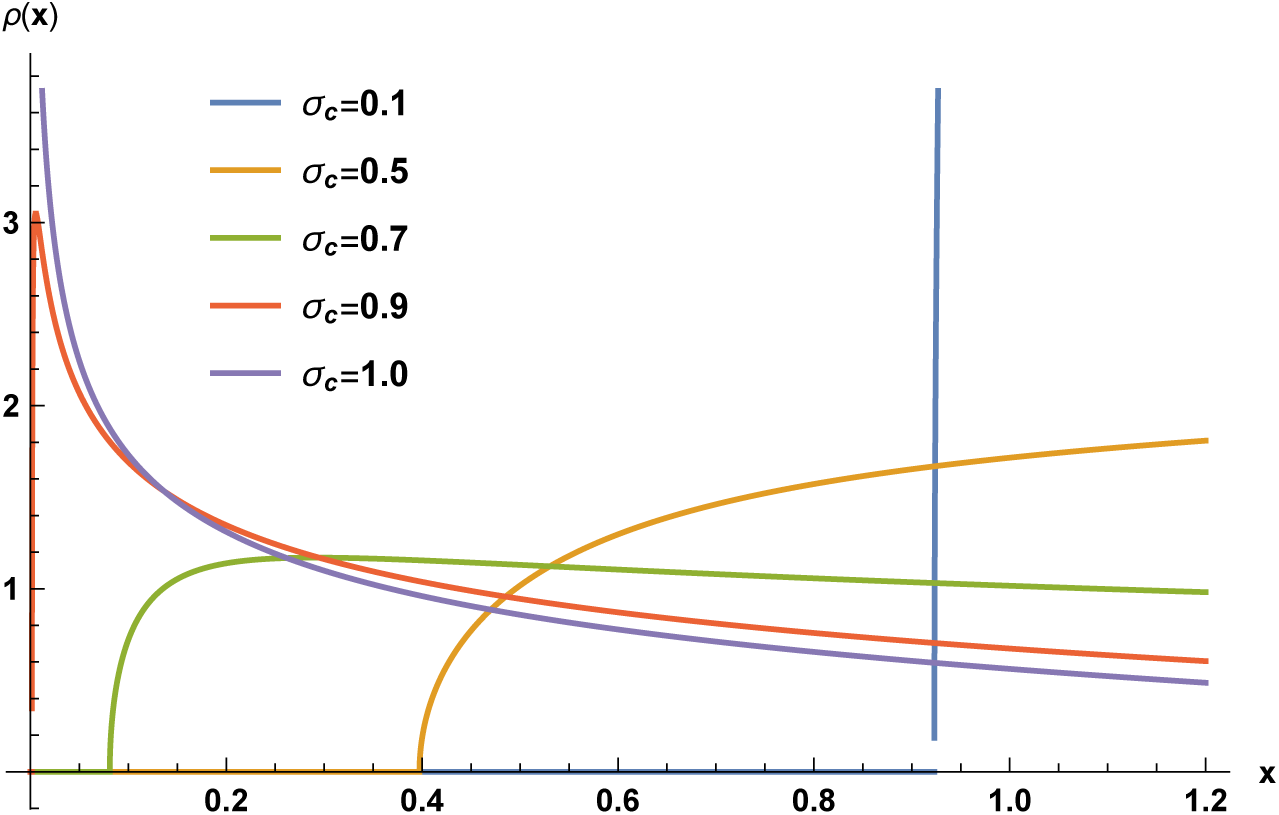
The asymptotic spectrum of *A*_*ij*_ for different values of *σ*_*c*_ by solving equation (43) numerically.

